# A novel allele of *SIR2* reveals a heritable intermediate state of gene silencing

**DOI:** 10.1101/2020.12.11.422089

**Authors:** Delaney Farris, Daniel S Saxton, Jasper Rine

## Abstract

Genetic information acquires additional meaning through epigenetic regulation, the process by which genetically identical cells can exhibit heritable differences in gene expression and phenotype. Inheritance of epigenetic information is a critical step in maintaining cellular identity and organismal health. In *Saccharomyces cerevisiae*, one form of epigenetic regulation is the transcriptional silencing of two mating-type loci, *HML* and *HMR*, by the SIR-protein complex. To focus on the epigenetic dimension of this gene regulation, we conducted a forward mutagenesis screen to identify mutants exhibiting an epigenetic or metastable silencing defect. We utilized fluorescent reporters at *HML* and *HMR*, and screened yeast colonies for epigenetic silencing defects. We uncovered numerous independent *sir1* alleles, a gene known to be required for stable epigenetic inheritance. More interestingly, we recovered a missense mutation within *SIR2*, which encodes a highly conserved histone deacetylase. In contrast to *sir1*Δ, which exhibits states that are either fully silenced or fully expressed, this *sir2* allele exhibited heritable states that were either fully silenced or expressed at an intermediate level. The heritable nature of this unique silencing defect was influenced by, but not completely dependent on, changes in rDNA copy number. Therefore, this study revealed a heritable state of intermediate silencing and linked this state to a central silencing factor, Sir2.

## Introduction

Transcriptional gene silencing is critical for proper cellular function, differentiation, and development. A temporally coordinated program of changing chromatin environments maintains cell fate by altering gene expression. Consequently, aberrant gene silencing and expression can lead to a variety of disease states (reviewed in Lee and Young 2013). A better understanding of how transcriptional silencing is maintained over time and remembered through cellular division is therefore crucial to understanding its misregulation.

One context in which transcriptional silencing has been studied in detail is the single-celled eukaryote *Saccharomyces cerevisiae. S. cerevisiae* exhibits stable, epigenetic silencing of transcription through the action of the SIR complex, which produces heterochromatin-like repressive chromatin domains (Kueng *et al*. 2013; Gartenberg and Smith 2016). This budding yeast has two mating types, *a* and *α*, with mating-type-specific information expressed from the two alleles of the *MAT* locus on Chromosome III. Two loci that undergo stable silencing are the silent mating type loci, *HML* and *HMR*. These extra copies of mating-type information are distal to the expressed *MAT* locus and allow for mating-type switches in the subset of strains with the *HO* gene, which encodes a site-specific nuclease that cuts at *MAT*.

*HML* and *HMR* are stably repressed, making mating type solely dependent on the allele of the *MAT* locus. Mutations in *SIR2, SIR3*, or *SIR4*, which collectively encode the SIR complex, result in complete loss of silencing at *HML* and *HMR* (Rine and Herskowitz 1987). The SIR complex is recruited to silencer elements within *HML* and *HMR*, deacetylates histones via the catalytic activity of Sir2, and binds to nucleosomes throughout the locus, resulting in transcriptional repression (Hoppe *et al*. 2002; Rusché *et al*. 2002; Thurtle and Rine 2014). Though Sir2, Sir3 and Sir4 are necessary for *HML* and *HMR* silencing, Sir1 was identified by mutant alleles that produced only partial loss of silencing at these loci (Rine *et al*. 1979). Characterization of the *sir1* phenotype at the single-cell level revealed that the expression states of *HML* and *HMR* are bistable in the absence of Sir1 (Pillus and Rine 1989). Quantitative RNA FISH studies show that in the silenced fraction of a *sir1Δ* population, *HML* and *HMR* are as fully silenced as in *SIR* cells (Dodson and Rine 2015). Likewise, in the unsilenced fraction, *HML* and *HMR* are as expressed as in *sir2*Δ, *sir3*Δ or *sir4*Δ mutants. These two expression states in *sir1*Δ are also heritable, as the mother cell’s expression state can be passed on faithfully to daughter cells for multiple generations, with switches to the opposite expression state occurring at a low rate.

In *sir1*Δ mutants, some cells manage to heritably silence *HML* and *HMR*, while others exhibit derepression of these loci. One possible explanation for the partial loss of silencing would be the existence of another gene with an overlapping function with *SIR1;* the absence of both factors would then be necessary to observe full derepression. Screens for enhancers of the *sir1*Δ silencing defect have largely uncovered more alleles of *SIR2, SIR3* and *SIR4* (Stone *et al*. 2000). Screens for suppressors of the silencing defect of *sir1*Δ mutants recovered *HTZ1*, which encodes a variant of histone H2A, and *ESC2* (Dhillon and Kamakaka 2000). However unlike *sir1*Δ, *htz1* and *esc2* do not exhibit a bistable phenotype. Therefore, mutants that function similarly to *sir1*Δ have eluded previous studies.

A screen to identify bistable silencing mutants has not previously been reported, nor have any reports appeared of heritable intermediate levels of gene silencing. In this study, we carried out a forward mutagenesis screen to identify metastable silencing mutants in *S. cerevisiae*. This screen differed from past screens in the use of fluorescent reporter genes at *HML* and *HMR*, providing the opportunity to observe silencing and heritability quantitatively at both the population and single-cell level.

## Materials and Methods

### Strains and Culture methods

All strains were derived from W303 and are listed in the Supplemental Material, Table S1. Plasmids used in the study are listed in Table S2. All oligonucleotides, used for cloning, PCR, and sequencing, are listed in Table S3. Strains were grown in Yeast Peptone Dextrose (YPD), or Complete Supplement Mixture (CSM) with or without individual amino acids left out (Sunrise Science Products), as indicated. The FLuorescent Analysis of Metastable Expression (FLAME) reporter strain was initially published in (Saxton and Rine 2019). Throughout this study, there were subtle differences between the silencing levels of *HML* versus *HMR;* however, the expression phenotypes of both remained similar. Elucidating any differences between the two loci was not pursued further. Mutagenesis was induced with Ethyl Methane Sulfonate (EMS). Diploid strains were created by genetic crosses and phenotypes were confirmed following sporulation by tetrad analysis. The point mutations within *SIR1* and *SIR2* were identified by PCR amplification and sequencing. Each mutant generated by mutagenesis was expected to contain multiple base-pair substitutions. Strains with single point mutations in the genes of interest were then engineered using Cas9 technology, as previously described (Lee *et al*. 2015; Brothers and Rine 2019). Single guide RNAs (sgRNAs) targeting *SIR2* and a unique linker region in *sir1Δ::LEU2* (JRY12861) are listed in Table S3. To generate *sir1* alleles in an unmutagenized parent strain, PCR-amplified repair templates of sequence-confirmed *sir1* alleles replaced the *sir1*Δ::*LEU2* allele in JRY12861. To create *sir2-G436D* in an unmutagenized parent strain, *sir2-G436D* was PCR amplified from JRY12466 and provided as a repair template to replace the *SIR2* allele in JRY12860. Both *sir1* and *sir2-G436D* mutant allele integration replaced the Cas9-directed cut site. All single point mutations were sequence confirmed. The Sir2-3xV5 fusion protein used for immunoblotting was created as described (Longtine *et al*. 1998). Strains were transformed with an amplified fragment of pJR3190 (Bähler *et al*. 1998), which allowed for homologous recombination and integration of the *KanMX* cassette and the 3x-V5 tag to the carboxyl terminus of the *SIR2* open reading frame. *SIR2* and *sir2-G436D* were amplified from JRY12860 and JRY12564, respectively, with 300 base pairs of 5’ promoter sequence and 200 base pairs of 3’ terminator sequence. These fragments were integrated into the 2 micron plasmid vector pRS426 to generate pJR3523 and pJR3524. Finally, *fob1*Δ::*KanMX* was generated by amplification of *KanMX* from pJR3190 and subsequent transformation.

### EMS mutagenesis

The EMS protocol was adapted from previously reported protocols (Winston 2008; Liu and Hu 2010) and optimized for our reporter strain (JRY12860) and reagents to yield ~50-60% lethality. Cells were plated at a low density (~150-200 colonies) on YPD for screening. Twelve independent rounds of mutagenesis were conducted. Approximately 11,000 mutagenized colonies were screened using fluorescence microscopy in the first eight rounds, and six mutants of interest were recovered. The final four rounds were initially screened in parallel via Fluorescent Activated Cell Sorting (FACS), as described below, and three mutants of interest were isolated (each mutant was selected from an independent culture). Mutants of interest were assigned a unique identifier during screening and identification (Figure S1), but throughout the text are referenced by the associated mutant allele, i.e. *sir1-P23S* or *sir2-G436D*.

### FACS Single Sort

Following EMS mutagenesis, independent mutagenized cultures were grown in liquid medium prior to parallel FACS sorting. Specifically, after the final resuspension in 500 uL YPD (~2 × 10^8^ cells), for each independent culture, 50 mLs of YPD were inoculated with the mutagenized cells and grown to saturation overnight. The following day, saturated cultures were back diluted to 0.1 OD in YPD and grown to log phase (~0.6-1.0 OD). 2 mL of each log-phase culture was harvested, washed, and resuspended in 2 mL 1X sterile PBS. Samples were strained through a 5 μm sterile mesh cap into a 5 mL polypropylene tube (Falcon), and kept on ice until sorting. *SIR+* and *sir4*Δ control strains (JRY12860, JRY12862) were used to determine fluorescence threshold levels, as all *SIR* cells were nonfluorescent, and all *sir4*Δ cells were fluorescent. One large gate for fluorescent cells (only GFP^+^, only RFP^+^, and GFP^+^ and RFP^+^) was created, and fluorescent cells (approximately 10,000 per culture) were sorted, grown at 30°C overnight, and then plated for screening. Plates were incubated at 30°C until colonies formed and were large enough for screening (2-4 days). Only a single mutant of interest was followed from each independent mutagenized culture.

### FACS Double sort

Double FACS sorting was performed for one round of mutagenesis. Strain JRY11906 was mutagenized. The first part of the double sort strategy was identical to the single sort strategy. Sorted fluorescent cells were then grown at 30°C in liquid culture overnight. Fresh YPD was inoculated with the daughters of the sorted cells to a density of 0.1 OD, and maintained near log-phase growth through continuous back dilution for two days, providing ample time for some fluorescent cells to switch into a silenced state. After these two days of growth, samples were prepared for sorting as above, but this time a gate for GFP^-^ and RFP^-^ cells was created, and these sorted cells were grown at 30°C in YPD overnight and prepared for colony screening as indicated above. Again, a single mutant of interest was followed per independent mutagenized culture.

### Colony Imaging

Cells were plated at a low density (20-35 cells/plate) on solid medium as indicated in individual figures. Single cells were then grown into colonies for 3-5 days at 30°C and imaged using a Leica M205 FA fluorescence stereo microscope and a Leica DFC3000 G microscope camera equipped with LAS X software (Leica). For all colony images within a given experiment, all conditions (growth, media, magnification, and exposure) were identical.

### Flow cytometry

Cells were inoculated into 150 uL of CSM in 96-well plates (Corning). Three biological replicates per strain were grown overnight. Saturated cultures were back diluted to ~0.1 OD in fresh CSM, and continuously back diluted to maintain log-phase growth for 24 hours; this growth period allowed the distribution of cells with silenced or non-silenced *HML* and *HMR* to reach equilibrium. After 24 hours of log-phase growth, cells were harvested by centrifugation, resuspended in 100 μL of 4% paraformaldehyde and incubated at room temperature for 15 minutes. Samples were pelleted and the fixed cells were resuspended in 100-150 uL of a 1X PBS solution. These fixed samples were stored at 4°C and analyzed by flow cytometry within 5 hours of fixation. Flow cytometry was performed using a BD LSRFortessa (BD Biosciences) with a FITC filter (for GFP) and a PE-TexasRed filter (for RFP), and at least 10,000 cells were analyzed per sample. Flow data was analyzed and visualized using FlowJo (BD Biosciences). All flow cytometry data were gated identically, omitting aggregates and cellular debris from analysis.

### Immunoblotting

Protein isolation, immunoblotting, and quantification were carried out as previously described (Brothers and Rine 2019). The membranes were blocked in Odyssey Blocking Buffer (LI-COR Biosciences), and the following primary antibodies and dilutions were used for detection: mouse anti-V5 (Invitrogen R960-25, 1:5,000) and rabbit anti-Hxk2 (Rockland #100-4159, 1:10,000). The secondary antibodies used were goat anti-rabbit (Li-Cor #C60531-05 1:20,000) and goat anti-mouse (Li-Cor #C81106-03 1:20,000). The immunoblot was imaged using a Li-Cor infrared fluorescent scanner.

### Patch mating assay

Strains to be assayed were patched onto solid YPD and grown at 30°C. After 1 day, these YPD plates were replica plated onto a mating-type tester lawn with complementary auxotrophic markers (*MATa* JRY2726, and *MATα JRY2728*, plated on YPD), and grown at 30°C for 1 day. Lawns were then replicated onto minimal YM medium, grown at 30°C for 2 days, and imaged.

### *α*-factor Confrontation Assay

This assay was carried out as previously described (Pillus and Rine 1989). Single, unbudded cells were micromanipulated approximately ½ field of view at 200X magnification away from the streak of *MATα* cells which served as a source of α-factor. The plates were incubated at 30°C for approximately 3 hours and the morphology of single cells was observed.

### Live-cell imaging

Strains JRY12861, JRY12564, and 12901 were grown as described above for flow cytometry, but in 5 mL cultures of CSM. After 24 hours of log-phase growth, a 500 μL aliquot of cell suspension at approximately 0.6-1.0 OD was harvested and resuspended in 500 uL sterile water. This cellular suspension was then sonicated for 5 seconds at 20% amplitude (Branson Ultrasonics Digital Sonifier 100-132-888R with Sonicator Tip 101-135-066R) to disrupt aggregates. A 5 μL aliquot of sonicated cells was spotted onto a CSM 2% agar pad. Once dry, the agar pads were inverted onto a 35 mm glass bottom dish (Thermo Scientific) and imaged using a Zeiss Z1 inverted fluorescence microscope with a Prime 95B sCMOS camera (Teledyne Photometrics), Plan-Apochromat 63x/1.40 oil immersion objective (Zeiss) filters, MS-2000 XYZ automated stage (Applied Scientific Instrumentation), and Micro-Manager imaging software (Open Imaging). Samples were incubated at 30°C and imaged every 10 minutes or 15 minutes for a total of 10 hours in bright-field, GFP, and RFP. Time-lapse movies were prepared and analyzed using FIJI software (Schindelin *et al*. 2012).

### Single-cell segmentation and fluorescence quantification

Bright-field microscopy images from live-cell imaging were segmented using the online tool Yeast Spotter (Lu *et al*. 2019) or manually for individual cells over long time courses, such as those shown in Figure 3B and C. Individual cells were parsed and labeled using the “analyze particles” tool in FIJI, and measurements taken, including area and GFP mean fluorescence intensity.

**Figure 1.**
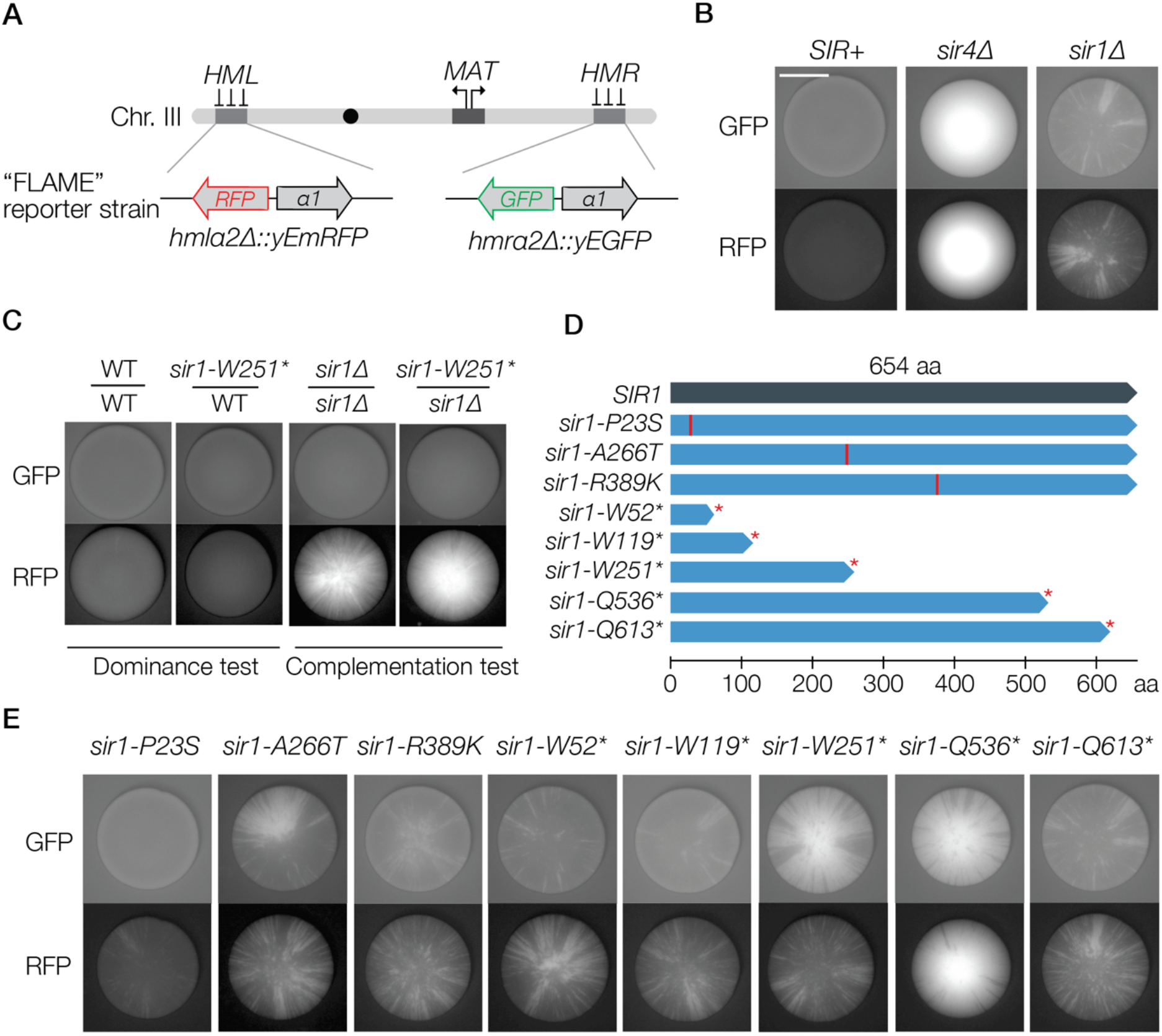
A screen for metastable silencing mutants revealed eight unique alleles of *sir1*. A. Schematic of the FLuorescent Analysis of Metastable Expression “FLAME” reporter strain (JRY12860) used in this study: fluorescent reporters *yEGFP* and *yEmRFP* replaced *α2* at *HMRα* and *HMLα*, respectively. B. Colony phenotypes of control strains (JRY12860, JRY1862, JRY1961) in both the GFP and RFP channel. Colonies were grown on YPD and imaged at identical exposures (Scale bar, 4 mm). C. Representative colony images of diploid strains for the dominance and complementation tests. For dominance testing, a *MATa* wild-type FLAME strain was mated with *MATα* mutant strain (JRY11955, JRY11915); for complementation testing, a *MATa sir1*Δ strain was mated with a *MATα* mutant strain (JRY11957, JRY11950). D. A schematic of the *sir1* alleles identified. The *SIR1* encodes a 654 amino acid protein (top bar in dark blue). Mutant alleles contained either a missense mutation or a nonsense mutation. Premature stop codons are indicated with an asterisk, i.e. *sir1-W52*\*. E. Representative colony images of the engineered single point mutation *sir1* alleles, imaged on YPD in both the GFP and RFP channel.

**Figure 2.**
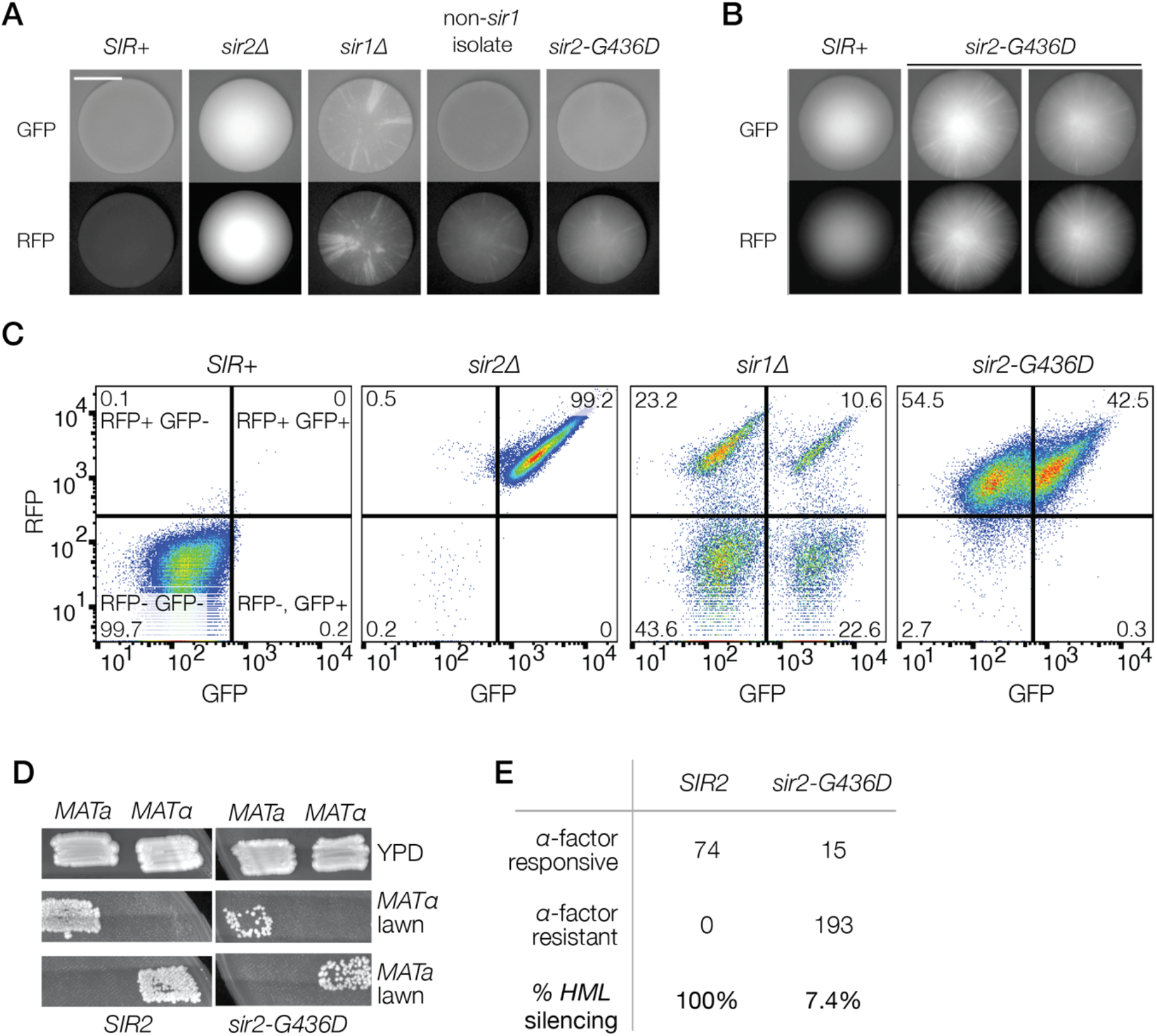
Characterization of mutant *sir2-G436D*. A. Representative colony images of FLAME control strains, the mutant of interest, and *sir2-G436D* in both the GFP and RFP channel (JRY12860, JRY12259, JRY12861, JRY12466, JRY12564), grown on YPD. B. Colony images of FLAME strain *SIR*+ colonies and two biological replicates of the engineered single point mutation strain (JRY12860, JRY12564). Colonies were grown on CSM and imaged at approximately 10-fold longer exposure than (A) (Scale bar, 4 mm). C. Flow cytometry plots of the fluorescence profiles for both *hmlα2*Δ::*RFP* (PE-Texas Red) and *hmrα2*Δ::*GFP* (FITC). Cells were grown in CSM liquid media for 24 hours, fixed, and analyzed. Quadrants were established using *SIR+* and *sir2*Δ strains (JRY12860, JRY12466), and the resulting percentage of the population per quadrant was labeled in the corresponding corner. *sir1*Δ cells (JRY12861) exhibited distinct populations in all four quadrants, while *sir2-G436D* (JRY12564) cells exhibited fully silenced states and intermediate silenced states. D. Patch mating assays of *SIR2* and *sir2-G436D* in *MATa* (JRY4012, JRY12667) and *MATα* (JRY4013, JRY12669) cells. The extent of growth on the YM minimal media reflected the strength of silencing. E. Results of the α-factor confrontation assay (JRY4012, JRY12667). *HML* silencing was calculated by dividing the number of α-factor responsive cells by the total number of cells assayed.

**Figure 3.**
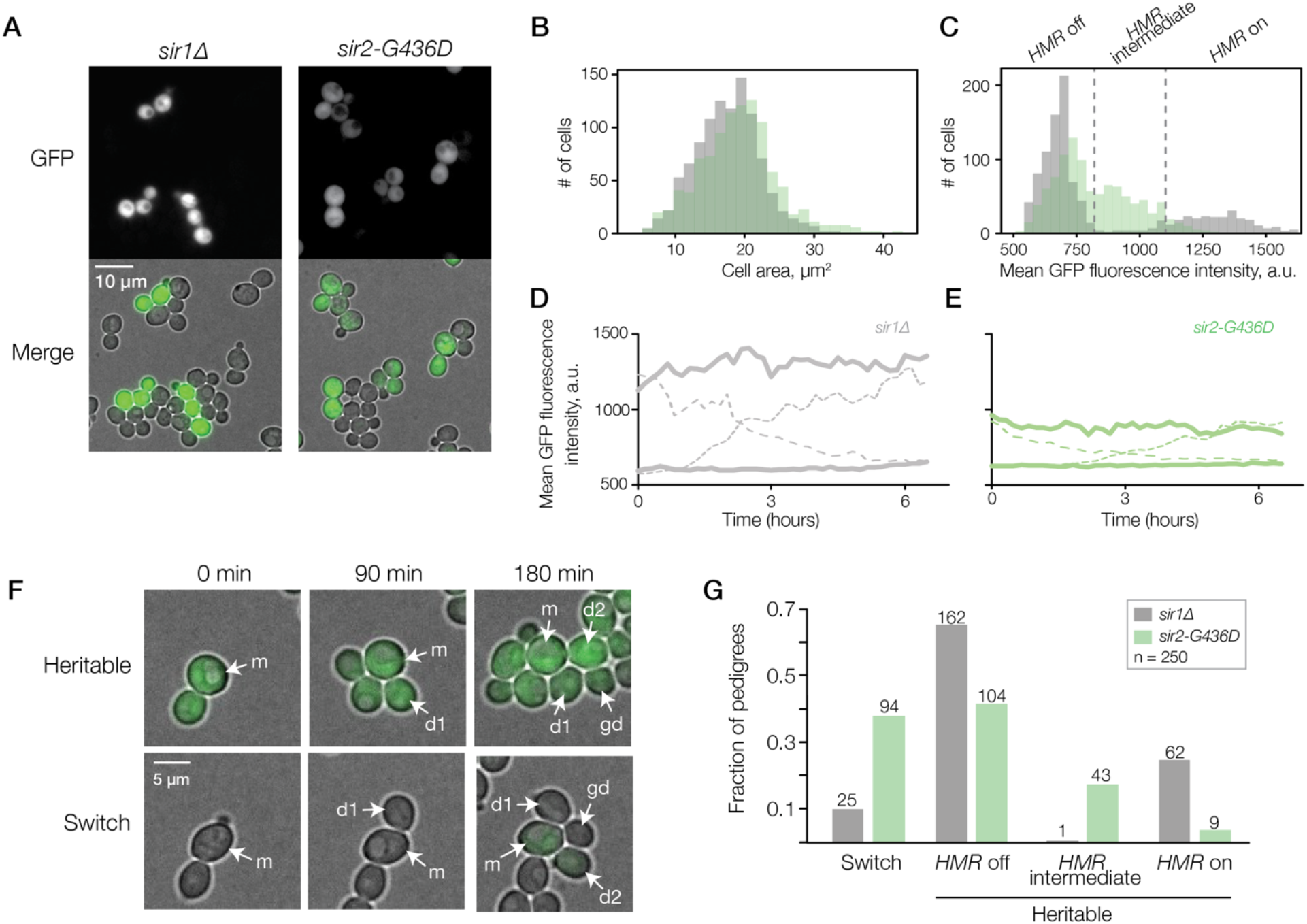
Live-cell imaging revealed the intermediate and heritable *sir2-G436D* expression state. A. GFP and merged (bright-field and GFP) fluorescence microscopy images of *sir1*Δ and *sir2-G436D* cells (JRY12861, JRY12564). B. Distribution of the cell size for both *sir1*Δ and *sir2-G436D*, with number of cells on the y-axis and cell area in μm^2^ on the x-axis. C. Distribution of the GFP mean fluorescence intensity (arbitrary units, a.u.) per cell for both *sir1*Δ and *sir2-G436D*. Dashed lines demarcate the boundaries of the three fluorescence states: *HMR* off, *HMR* intermediate, and *HMR* on. Details on how thresholds were assigned are in the *Methods*. D. GFP mean fluorescence intensity for individual *sir1*Δ cells over 6.5 hours. 12 individual cells were monitored, and representative fluorescence trajectories are displayed. Each solid line represents a single cell that maintained a similar fluorescence level over the timecourse, whereas each dashed line represents a single cell that experienced a change in fluorescence. E. Same as (D), but for individual *sir2-G436D* cells. F. An example of the pattern of divisions and pedigrees designated as “heritable” versus a “switch” in *sir2-G436D* cells. A single mother cell (m, t = 0 min) budded twice, producing daughter 1 (d1, t = 90 minutes) and daughter 2 (d2, t = 180 minutes). Budding of daughter 1 gave rise to a grand-daughter (gd, t = 180 minutes) cell. In the “heritable” example, all cells at all time points displayed a fluorescence level falling within the “*HMR* intermediate” range; in the “switch” example, a loss of silencing occured during the second division, giving rise to cells with fluorescence classified as “*HMR* intermediate”. G. Bar chart showing the percentage of pedigrees designated as a “switch” or “heritable”. 250 pedigrees were observed per genotype, with the number of pedigrees per category above each bar.

Individual cells were assigned a silencing state of *hmrα2*Δ::*GFP* (“*HMR* off”, “*HMR* intermediate”, “*HMR* on”), using threshold values determined from the *sir1*Δ (JRY12861) single cell-analysis data. The *sir1*Δ cell data was split into two populations (“*sir1*Δ off” and *“sir1*Δ on”), with each population assumed to be normally distributed, and the threshold values designated to include 90% of the respective *sir1* population. The “*HMR* off” to “*HMR* intermediate” boundary was defined as the 90^th^ percentile rank GFP fluorescence intensity value for the “*sir1*Δ off” population (798 GFP mean fluorescence intensity). The “*HMR* intermediate” to “*HMR* on” boundary was defined as the 5^th^ percentile rank GFP fluorescence intensity value for the “*sir1*Δ on” population (1103 GFP mean fluorescence intensity).

### Pedigree analysis for measuring heritability

Time-lapse microscopy movies (described above) were analyzed to measure the heritability of a fluorescence state. For each pedigree analyzed, a single cell (mother) and the resulting three progeny (daughter 1, daughter 2, grand-daughter (daughter of daughter 1)) were manually segmented and the fluorescence state was measured using FIJI software. At each time point (t = 0 minutes, t = 90 minutes, t = 180 minutes), all cells were measured and assigned a fluorescence state, using the threshold values established above. If all progeny at all time points displayed the same expression state as the mother cell had at t = 0 minutes, the pedigree was labeled “heritable” reflecting the heritability of that expression state. If any of the cells in the pedigree switched state designations, the pedigree was labeled as a “switch”, reflecting the absence of heritability of that expression state in that pedigree.

## Results

### Identification of metastable mutants

To isolate mutants that displayed metastable silencing defects at *HML* and *HMR*, we utilized an assay that reveals the expression state of these two loci individually. The FLuorescent Analysis of Metastable Expression (FLAME) assay utilizes fluorescent reporters integrated at *HML* and *HMR*, termed *hmlα*2Δ::*RFP* and *hmrα*2Δ::*GFP*, respectively (Saxton and Rine 2019, Figure 1A). In wild-type cells, these loci are stably silenced by the SIR complex (Rine and Herskowitz 1987). Thus, when SIR complex members Sir2, Sir3, or Sir4 are absent, these loci are fully expressed. In the FLAME assay, loss of silencing results in expression of the fluorescence reporters, which can be evaluated at either the single-cell or colony level. The colony phenotype offers additional historical information about the expression state of *HML* and *HMR*. Due to the pattern of cell divisions, ancestors are proximal to their descendants, forming sectors of related cells which radiate to the periphery of the colony. In *sir2, sir3*, or *sir4* mutants, colonies are uniformly fluorescent, whereas in a *sir1*Δ mutant, a sectored fluorescence pattern is observed; this sectoring indicates heritable phenotypic variation within a genetically identical population (Figure 1B). By screening colonies arising from the mutagenized *SIR* reporter strain, we identified six mutants with metastable silencing of *HML* and *HMR* (Figure S1A).

As a complement to direct screening of colonies, we adapted Fluorescence Activated Cell Sorting (FACS) to detect and sort fluorescent cells within a mutagenized population. These sorted cells were then interrogated for clonal heritability of expression states at the colony level (see Methods). Using a double-FACS sorting strategy, three additional mutants of interest were found (Figure S1A).

### Genetic analysis identified eight unique *sir1* alleles

The metastable phenotype was recessive in all nine mutants of interest, based on the fluorescence of diploids heterozygous for the new mutations (Figure 1C and S1B). To test if the metastable phenotype was due to a single mutation, seven of the diploids from the dominance test were sporulated and the phenotype evaluated among the tetrad segregants. The characteristic 2:2 segregation of mutant to wild-type phenotypes was observed for at least ten tetrads from each of the mutants tested, strongly suggesting that a mutation in a single gene caused the metastable phenotype.

A complementation test was used to determine whether these mutants revealed new genes capable of metastable phenotypes or were new alleles of *SIR1*. In this test, *MATα* mutants were mated to an isogenic *MATa sir1*Δ strain. All seven mutants tested failed to complement a *sir1*Δ mutation (Figure 1C and S1C). Interestingly, in diploids, the silencing phenotype was less severe than in haploids, and far more evident at *HML*. This discrepancy likely reflects previous findings that silencing is stronger in diploids than in haploids (Dodson and Rine 2015), and that haploid *sir1*Δ cells are more frequently silenced at *HMR* than at *HML* (Saxton and Rine 2019).

The *sir1* alleles of each mutant strain were sequenced, revealing mutations within the coding region of *SIR1* (Figure 1D). Two independent rounds of mutagenesis produced identical nonsense mutations, resulting in identical *sir1-W251\** alleles. As expected from EMS mutagenesis, all of the *sir1* alleles contained a single point mutation resulting from GC to AT transitions, with five of the eight unique point mutations resulting in a nonsense mutation (Figure S1D). These point mutations were engineered into the parent strain using molecular cloning techniques, where they recapitulated the phenotypes observed in the original mutants, showing that the *sir1* alleles were necessary and sufficient to produce the metastable phenotype observed (Figure 1E).

### A metastable phenotype from a mutation in *SIR2*

Having identified eight independent and unique *SIR1* alleles, we revised the screening strategy to reduce the likelihood of finding more *sir1* mutants. We reasoned that if two *SIR1* alleles were present in our haploid reporter strain, the probability of random mutagenesis disrupting both in the same cell would be reduced. Therefore, an additional copy of *SIR1* was maintained on a plasmid in the parental strain of the screen (JRY12860 containing pJR909). After mutagenesis, a single-FACS enrichment step was employed (see Methods). With this additional extrachromosomal copy of *SIR1*, very few mutants with a metastable phenotype were produced. After mutagenizing and sorting twelve independent cultures with FACS, no further *sir1* alleles were found. One colony of interest was identified, which exhibited a mild but noticeable silencing defect (Figure 2A). The phenotype was unlike any other observed during both iterations of mutagenesis and unique from all other mutant phenotypes studied using the FLAME assay. In this mutant, the entire colony exhibited expression of *HML* and *HMR*, but the strength of expression was less than that observed in *sir2Δ*. Moreover, close examination of the colony revealed streaks of greater or lesser fluorescence intensity, suggesting the possibility of heritable intermediate defects in silencing, a phenotype not previously described. The mutant phenotype was recessive, complemented a *sir1*Δ mutation, and produced a 2 wild-type: 2 mutant segregation pattern after diploid sporulation and tetrad analysis (Figure S2).

To identify the causative gene resulting in the mutant phenotype, we first assayed the ability of *SIR2, SIR3*, or *SIR4* to rescue the silencing defect. Transformation of a *SIR2* plasmid into the parent strain restored wild-type silencing, whereas *SIR3* and *SIR4* plasmids had no effect on the silencing phenotype. Sir2 is a highly conserved histone deacetylase and is the sole catalytic component of the SIR complex (Landry *et al*. 2000; Imai *et al*. 2000). Sequencing of *SIR2* from the mutant strain revealed a single point mutation at residue 436, changing the encoded amino acid from a glycine to an aspartic acid (*sir2-G436D*). Using molecular cloning techniques, the *sir2-G436D* point mutation was introduced into the parental strain (JRY12564); this mutant recapitulated the intermediate silencing phenotype (Figure 2A). Thus, the missense *sir2-G436D* allele was sufficient to produce the intermediate silencing defect. Colony imaging at longer exposures highlighted the unique fluorescence pattern of this mutant, with streaks of brighter fluorescence superimposed on a low-fluorescence colony (Figure 2B). Interestingly, these streaks of brighter fluorescence overlapped in the RFP and GFP channels, suggesting that *hmlα2*Δ::*RFP* and *hmrα2*Δ::*GFP* were coordinately impacted by *sir2-G436D*. This result contrasted with the colony phenotype of *sir1*Δ, in which *hmlα2*Δ::*RFP* and *hmrα2*Δ::*GFP* are silenced or expressed independently of each other (Figure 2A, Xu *et al*. 2006).

### A unique silencing defect in *sir2-G436D*

To further characterize this mutant phenotype, flow cytometry was used to quantify the *hmlα2*Δ::*RFP* and *hmrα2*Δ::*GFP* fluorescence intensities of log-phase cells. The *SIR* reporter strain existed as a homogenous population lacking both GFP and RFP fluorescence, whereas the *sir2*Δ strain strongly expressed both GFP and RFP (Figure 2C). Using the *SIR+* and *sir2*Δ control strains, gates were established to create four quadrants representative of the four possible FLAME expression states. As expected, *sir1*Δ cells existed in all four quadrants and therefore exhibited all possible combinations of expression states for *HML* and *HMR*. The *sir2-G436D* mutant strain exhibited a distinct pattern of expression, with a broad spread in fluorescence intensities for *hmrα2*Δ::*GFP* and *hmlα2*Δ::*RFP*. The distribution of fluorescence intensities appeared bimodal for *hmrα2*Δ::*GFP*, and distinctly less so for *hmlα2*Δ::*RFP*, which was more uniformly expressed. Interestingly, GFP^+^ cells and RFP^+^ cells appeared less fluorescent in *sir2-G436D* than in *sir1*Δ or *sir2*Δ. Therefore, flow cytometry indicated that some *sir2-G436D* cells exhibited a fully silenced state, while others exhibited an intermediate silenced state.

To evaluate this intermediate silencing phenotype further, the *sir2-G436D* allele was introduced into a strain with wild-type *HML* and *HMR*. Using these strains, silencing of *HML* and *HMR* was measured by a patch mating assay and a single-cell α-factor confrontation assay. In the patch mating test, *sir2-G436D* silenced *HML* and *HMR* inefficiently relative to wild type (Figure 2D). An α-factor confrontation assay (Pillus and Rine 1989) revealed that approximately 7% of *MATa sir2-G436D* cells were able to sufficiently silence *HML* and thus avoid the the α-factor resistance of pseudo *a/α* diploids (Figure 2E). Compared to *sir1*, which by α-factor confrontation was previously shown to effectively repress *HML* in 20% of cells, *sir2-G436D* showed a more pronounced silencing defect. Thus, as confirmed by three independent assays, the *sir2-G436D* mutation resulted in partially defective silencing.

### *sir2-G436D* produced intermediate, heritable expression

To monitor the different silencing states of *sir1*Δ and *sir2-G436D* over time, we first tested whether these states were evident by live-cell microscopy. To simplify the analysis of the microscopy, we focused on the expression states of *hmrα2*Δ::*GFP*. As previously established by RNA FISH (Dodson and Rine 2015), *sir1*Δ cells exhibited either full silencing or full expression of *HMR* (Figure 3A). In partial contrast, *sir2-G436D* cells exhibited full silencing or partial silencing of *HMR* (Figure 3A). To quantify fluorescence levels, cell segmentation and quantification were performed. Using bright-field images, individual cells were segmented with Yeast Spotter (Lu *et al*. 2019). Once segmented, single-cell data were extracted and displayed as a histogram (Figure 3B and C). As anticipated, cell size was approximately normally distributed, with no meaningful difference between the genotypes (Figure 3B); however, the GFP fluorescence profiles per genotype were distinct. *sir1*Δ cells were either silenced or fully expressed, similar to the flow cytometry data, whereas *sir2-G436D* cells were either fully silenced or partially silenced (Figure 3C). Using fluorescence intensities of *sir1*Δ cells, threshold values were established to demarcate three fluorescence states: “*HMR* off”, “*HMR* intermediate”, and *“HMR* on” (Methods). Using these thresholds, approximately 40% of the *sir2-G436D* cells measured exhibited intermediate fluorescence (“*HMR* intermediate”), while only 4% of *sir1*Δ cells displayed intermediate expression.

For a transcriptional state to be classified as epigenetic, it must be heritable through cell divisions. Therefore, we assessed the ability of the *sir1*Δ and *sir2-G436D* mutants to reliably transmit the observed silencing states over multiple generations. Time-lapse movies of dividing cells qualitatively suggested that the observed states were heritable (Supplemental Movies 1-2). To quantitatively assess this heritability, we monitored the fluorescence of individual mother cells over the course of four division events, or approximately six hours. In both *sir1*Δ and *sir2-G436D*, the fluorescence states of individual cells could be maintained over this entire period or switch to a different state (Figure 3D and E). To calculate approximate switching rates, unbudded cells and the resulting progeny of two generations were manually tracked, with each cell assigned an *HMR* expression state according to the threshold values in Figure 3C. A pedigree was designated as “heritable” if all cells within the pedigree exhibited the same expression state of *HMR* at all three time points (Figure 3F). In contrast, the pedigree was designated as a “switch” if any of the cells within the pedigree switched to a different *HMR* expression state. Two generations were analyzed to increase our confidence that the *HMR* expression states were heritable and did not simply reflect variation in fluorescent properties of individual cells. Using this method, 250 pedigrees per genotype were analyzed (Figure 3G).

Recent studies using microscopy and flow cytometry show that approximately 10% of *sir1*Δ cell divisions give rise to a switch in *HMR* expression state (Saxton and Rine 2019). Consistent with this finding, approximately 10% of *sir1*Δ pedigrees analyzed resulted in a switch in *HMR* silencing, while the other 90% of pedigrees displayed heritability (Figure 3G). The occurrence of switching in *sir2-G436D* was higher than in *sir1Δ*. However, a majority of pedigrees, approximately 62%, displayed heritability of the *HMR* expression state. Though the majority of these heritable pedigrees displayed “*HMR* off” silencing, 28% of the *sir2-G436D* pedigrees analyzed showed stable transmission of the “*HMR* intermediate” state. This live-cell imaging analysis further supported that *sir2-G436D* exhibited an intermediate silenced state and showed that this state was inherited through cellular division.

### *sir2-G436D* silencing defects were partially due to reduced levels of Sir2

Based on the crystal structure of the Sir2 protein (Hall and Ellenberger 2008; Hsu *et al*. 2013), codon 436 falls within the highly conserved C-terminal catalytic domain. However, residue 436 is distinct from the site of catalysis and is in close proximity to the zinc ion within the zinc-finger domain (Figure 4A). A previous study found that disruption of the zinc-finger domain by mutation of the coordinating cysteine residues results in full silencing loss (Sherman *et al*. 1999). Strikingly, the aspartic acid introduced by the *sir2-G436D* mutation is predicted to encroach on the zinc-coordinating site, which may disrupt the protein stability and silencing capacity of Sir2-G436D (Figure 4A).

**Figure 4:**
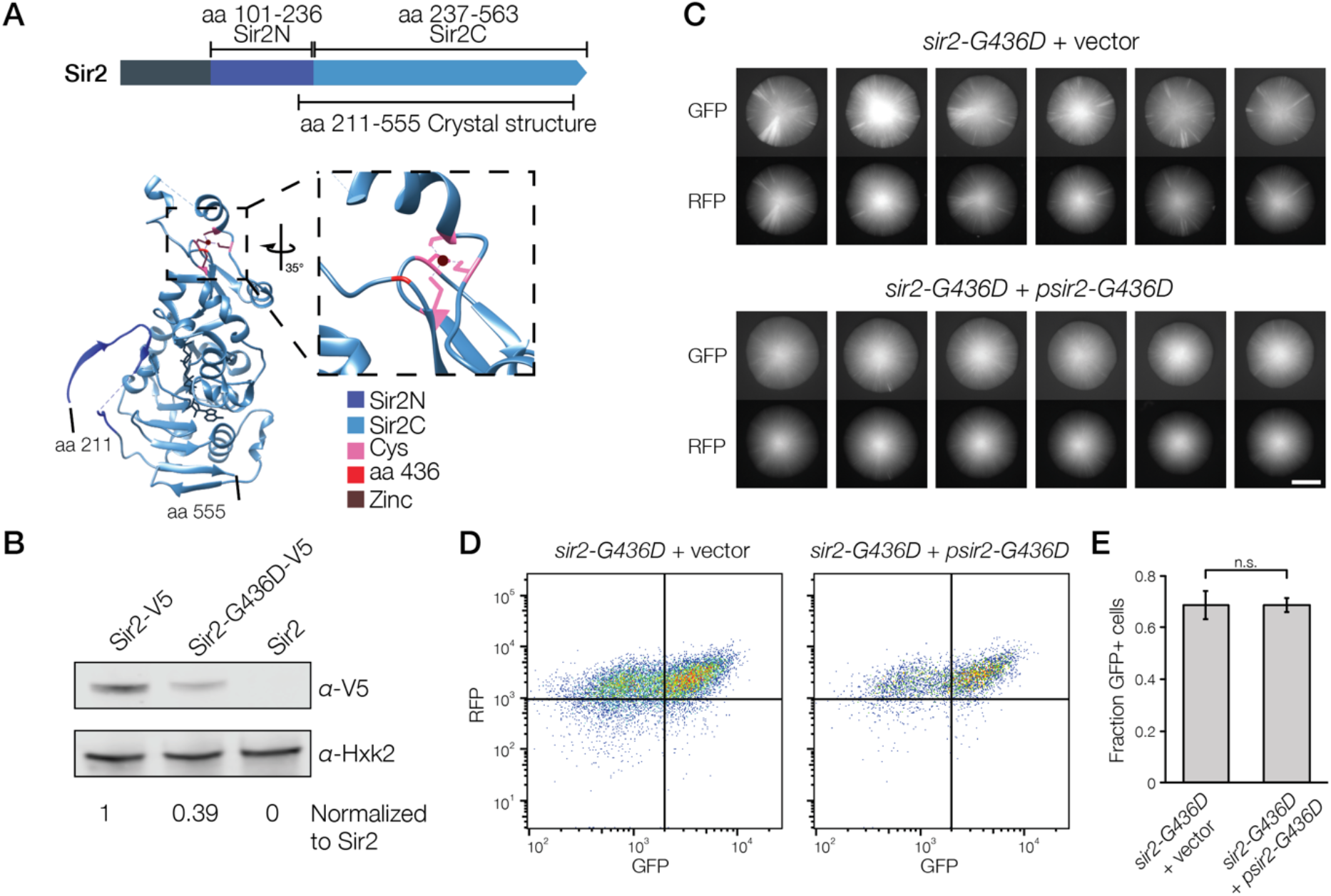
Sir2-G436D levels were partially responsible for variegated silencing. A. A schematic of the Sir2 protein and its crystal structure (Hall and Ellenberger 2008, Hsu *et al*. 2013). The N-terminal helical domain (dark blue) and C-terminal catalytic domain (light blue) are indicated. The crystal structure spans from amino acid 211-555 and contains a zinc ion (brown), zinc-coordinating cysteines (pink), and the site of the Sir2-G436D point mutation (red). The inset shows the zinc-coordinating site in Sir2. B. Immunoblot to detect Sir2-V5, Sir2-G436D-V5, and an internal loading control Hxk2 (JRY12589, JRY12590). Protein levels were quantified, normalized to the loading control, and compared to wild-type Sir2-V5 levels. C. Representative colony images of *sir2-G436D* (JRY12564) plus a 2 micron plasmid vector (pRS426) or a 2 micron plasmid containing *sir2-G436D* (pJR3525). Six colonies are shown per genotype. Colonies were grown on CSM∓Ura to select for plasmids. Scale bar, 3 mm. D. Representative flow cytometry profiles of same strains shown in (C). Independent cultures (n = 3 per genotype) were grown at log phase for 24 hours in CSM −Ura liquid media, fixed, and analyzed. Representative flow cytometry profiles for each strain are shown. Quadrants were established by using the fluorescence profiles of *SIR2* and *sir2*Δ cells (Figure S3). E. Fraction of GFP^+^ cells in independent cultures grown in (D). Data are means ± SD (n = 3 independent cultures per genotype). A two-tailed t-test was used for statistical analysis.

To test whether this mutation affected the stability of Sir2, the wild-type and mutant Sir2 proteins were tagged with the V5 epitope and protein levels were evaluated by immuno-blot. Mutant Sir2-G436D levels were roughly 40% of the wild-type Sir2 levels (Figure 4B). If this reduced expression was responsible for the observed silencing defects, we would expect that overexpression of *sir2-G436D* would ameliorate these defects. Indeed, expression of *sir2-G436D* from a high copy number plasmid reduced the amount of variegation in the *sir2-G436D* mutant strain, as compared with a vector-only control (Figure 4C, Figure S3A). These data suggested that the *sir2-G436D* silencing defect was partially due to reduced levels of Sir2-G436D. Surprisingly, the effects of this *sir2-G436D* plasmid were not observed by flow cytometry (Figure 4D and E, Figure S3B). This discrepancy provided an early indication that the variegation observed in *sir2-G436D* colonies is a relatively small part of the heritability observed at the single-cell level. This idea is explored further in the subsequent section.

### rDNA recombination accounted for variegated silencing in *sir2-G436D* colonies

In addition to its role in silencing at *HML, HMR*, and telomeres, Sir2 is also part of the RENT complex, which binds to rDNA and suppresses recombination between rDNA repeats (Gottlieb and Esposito 1989; Straight *et al*. 1999; Kobayashi *et al*. 2004). Though this activity stabilizes the rDNA copy number, the copy number can still expand and contract in *SIR2* cells. A previous study found that cells with low rDNA copy numbers exhibit stronger heterochromatic silencing at an artificial telomere and destabilized version of *HMR*, suggesting that the SIR complex competes with the RENT complex for a limiting amount of Sir2 (Michel *et al*. 2005). By extension, this study suggests that different rDNA copy numbers require different amounts of the RENT complex, which changes the amount of Sir2 that is available for heterochromatic silencing. Therefore, heritable differences in rDNA copy number may lead to the heritable differences in silencing efficiency in *sir2-G436D*.

Fob1 is a nucleolar protein that functions to create replication fork barriers in the rDNA, which prevent collisions between DNA polymerase and RNA polymerase I (Kobayashi and Horiuchi 1996; Kobayashi *et al*. 1998). Additionally, replication fork barriers generate recombinogenic replication intermediates that drive the expansion and contraction of rDNA repeats. Thus, in the absence of *FOB1*, recombination in the rDNA is greatly reduced. To test if changes in rDNA copy number contributed to changes in silencing states of *HML* and *HMR*, we generated a *sir2-G436D, fob1*Δ double mutant. In comparison to *sir2-G436D*, the *sir2-G436D, fob1*Δ double mutant exhibited substantially less variegation of *HML* and *HMR* expression at the colony level (Figure 5A, Figure S4A). These data suggested that rDNA recombination plays a role in the *sir2-G436D* silencing defect.

**Figure 5:**
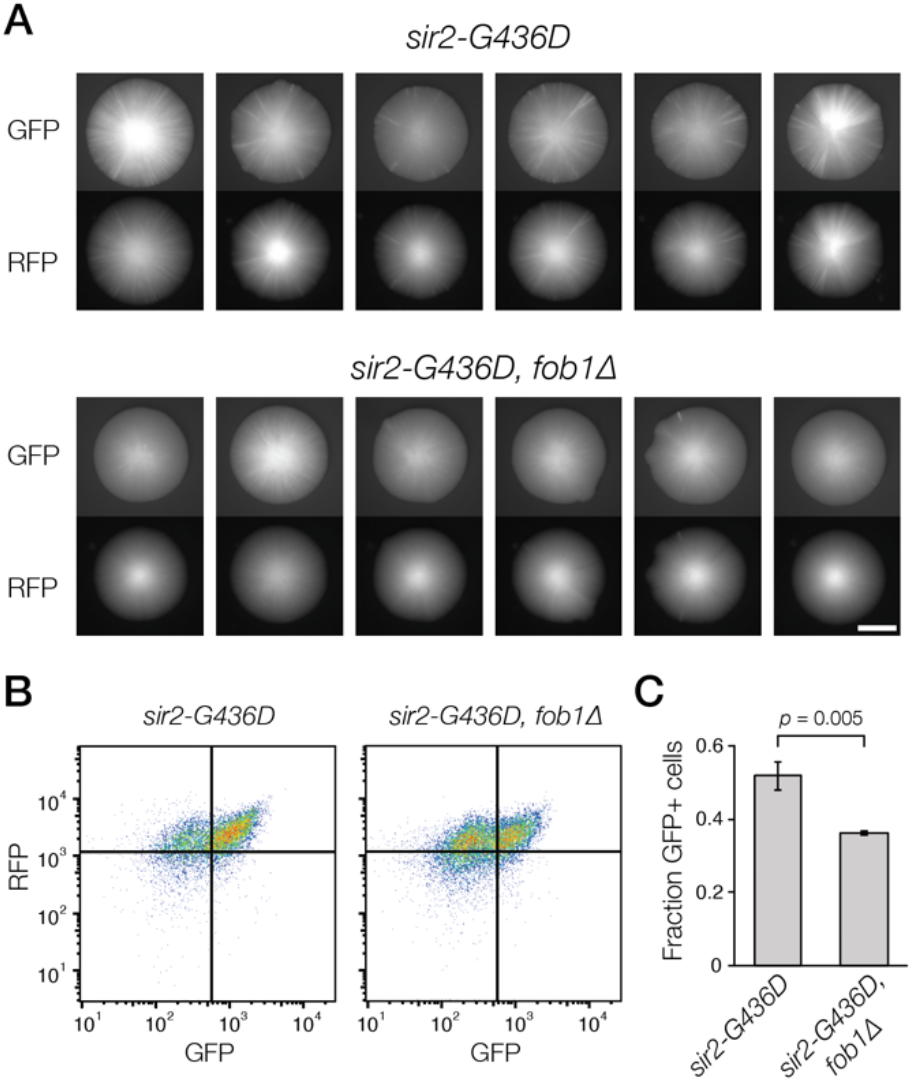
Changes in rDNA copy number were partially responsible for silencing variegation in *sir2-G436D*. A. Representative colony images of *sir2-G436D* (JRY12564) and the *sir2-G436D, fob1*Δ double mutant (JRY12901). Six colonies are shown per genotype. Colonies were grown on CSM. Scale bar, 3 mm. B. Flow cytometry profiles of same strains shown in (A). Independent cultures (n = 3 per genotype) were grown at log phase for 24 hours in CSM liquid media, fixed, and analyzed. A representative flow cytometry flow profile for each strain is shown. C. Fraction of GFP^+^ cells in independent cultures grown in (B). Data are means ± SD (n = 3 independent cultures per genotype). A two-tailed t-test was used for statistical analysis.

Though the variegation at the colony level was strongly reduced in the *sir2-G436D, fob1*Δ double mutant, these colonies still exhibited a uniform hazy fluorescence. Therefore, we tested whether *fob1*Δ altered the fluorescence profiles of single cells. Similar to the *sir2-G436D* single mutant, *sir2-G436D, fob1*Δ double mutant cells were either fully silenced or silenced to an intermediate level, and exhibited switching events between these two states (Figure 5B, Supplemental Movie 3). However, the double mutant consistently exhibited fewer cells in the intermediate silenced state (Figure 5B and C). These data suggested that the heritability of silencing at the single-cell level was partially independent of rDNA copy number. In this framework, our data indicated that *sir2-G436D* silencing defects reflected an admixture of two phenomena: (1) switching events that occured with a high frequency at the single-cell level, which manifested as intermediate, hazy fluorescence at the colony level and was not heavily influenced by rDNA copy number, and (2) switching events that were difficult to observe in single cells, but were readily observed at the macroscopic level of a colony, and due to changes in rDNA copy number.

## Discussion

The ability of cells to “remember” a silenced state has historically been uncovered by mutations that generate variegated expression. Despite the value of these mutations, such as *sir1*Δ, previous studies have not systematically screened for variegated silencing phenotypes in *S. cerevisiae*. Here, we performed a metastability screen that uncovered multiple new alleles of *SIR1*, but also identified a novel allele of *SIR2* that exhibited a heritable, intermediate silenced state. Further characterization of *sir2-G436D* revealed that the heritability of this state was not based on rDNA copy number, though changes in rDNA copy number influenced the silencing profile at the colony level.

### Sir1 was the main factor preventing metastable silencing of *HML* and *HMR*

Using a forward genetic screen and an assay for metastable silencing defects, we identified nine independent mutant alleles of *sir1*, of which eight were unique. Thus, to a first approximation, the screen had been saturated. It was therefore unlikely that variable penetrance of the *sir1*Δ silencing phenotype was due to a second non-essential gene with overlapping function. Once an additional copy of *SIR1* was introduced for screening purposes, no further *sir1* alleles were found, and very few mutants displayed a metastable phenotype. These results strongly suggested that Sir1 was the most important protein in converting silencing of *HML* and *HMR* from a metastable to fully silenced regime. This idea is consistent with a previous study in which metastable silencing at a telomeric reporter was strengthened by ectopic recruitment of Sir1 (Chien *et al*. 1993).

### The unique phenotype of *sir2-G436D*

A novel mutation, *sir2-G436D*, was identified with two striking qualities: 1) The mutation created an intermediate level of silencing, which was heritable through cell divisions as documented by single cell analysis. 2) At the colony level, this intermediate level of silencing was accompanied by radial streaks of cells with different expression states of the fluorescent reporters. Before discussing the phenotype of this mutant in detail, it is useful to consider the growth dynamics of a yeast colony. Any cell in a colony is a descendant from its more centrally located ancestors. When there is a heritable change in the expression state of a fluorescent reporter gene, that expression state is propagated outward, resulting in a wedge-shaped sector of cells that all exhibit the same state. Thus, a fluorescent sector represents a historical record of a transcriptional switching event that occurred at the apex of the sector, and that was inherited during subsequent colony growth.

The colony-level phenotype of *sir2-G436D* differed from that of *sir1*Δ in multiple ways. First, fluorescent sectors were less fluorescent in *sir2-G436D*, suggesting that the cells in these streaks also had an intermediate level of silencing. Second, the fluorescent sectors were more frequent in *sir2-G436D*, indicating that the switching rate between expression states differed from that seen in *sir1Δ*. Finally, *sir2-G436D* exhibited high concordance between the GFP and RFP channel (Figure 2B), implying that *HML* and *HMR* were coordinately impacted during the majority of the colony growth. This observation strongly suggested that the process responsible for radial streaks of fluorescence acted in *trans*. In contrast, the expression states of *HML* and *HMR* behave independently of each other in *sir1*Δ ((Xu *et al*. 2006), Figure 1B), demonstrating *cis*-transmission of expression states in this context. Together, these data suggested that the variegated expression seen in *sir2-G436D* and *sir1*Δ colonies were driven by fundamentally different mechanisms.

### rDNA copy number contributed to variegated expression in *sir2-G436D*

Given that deletion of *SIR2* causes full loss of silencing, it was likely that *sir2-G436D* was a hypomorphic allele. The G436D mutation was predicted to affect the zinc finger domain by generating a large polar side chain that disrupted the zinc finger domain (Figure 4). A previous study found that mutation of the four cysteine residues that coordinate with the zinc ion does not affect Sir2 levels but abolishes the silencing capacity of this protein (Sherman *et al*. 1999). In contrast, Sir2-G436D protein levels were reduced by 40% compared to wild-type Sir2 and exhibited a partial silencing defect. This dichotomy suggested that the Sir2-G436D may have partially disrupted the function of the zinc-coordinating domain and destabilized the mutant protein. Thus, altered levels of Sir2-G436D may be responsible for the silencing defects observed in this mutant. Consistent with this idea, overexpression of *sir2-G436D* from a high copy number plasmid strongly reduced silencing variegation observed at the colony level (Figure 4).

Sir2 is a protein that has multiple functions at different genomic locations. At silenced loci, Sir2 is part of the Sir2/3/4 complex and functions to deacetylate H4K16, which is necessary for silencing (Moazed and Johnson 1996; Landry *et al*. 2000; Imai *et al*. 2000). Separately, Sir2 is part of the RENT complex at rDNA repeats, where it stabilizes rDNA copy number by repressing transcription and regulating cohesin dynamics (Gottlieb and Esposito 1989; Straight *et al*. 1999; Kobayashi and Ganley 2005). Previous studies demonstrate that lower rDNA copy numbers enhance Sir2/3/4-dependent silencing at telomeres, suggesting that the RENT complex and Sir2/3/4 complex compete for a limited amount of Sir2 (Michel *et al*. 2005). We hypothesized that this competition for Sir2 was the underlying mechanism for the variegation observed in *sir2-G436D*. In this model, variation in rDNA copy number would change the amount of rDNA-bound RENT complex, which would then change the amount of Sir2 available for silencing at loci such as *HML* and *HMR*. This model would be consistent with coregulation of *HML* and *HMR* observed at the colony level in *sir2-G436D*, as altered levels of free Sir2 would influence *HML* and *HMR* equally in *trans*.

This model predicted that cells with a reduced ability to change rDNA copy number would exhibit reduced variegation of silencing in *sir2-G436D*. Indeed, removal of *fob1*, which is necessary for rDNA recombination, strongly reduced the silencing variegation of *HML* and *HMR* in this context. These data strongly suggested that the heritability of expression states observed in *sir2-G436D* was due to rDNA copy number. In light of this finding, we speculated that under normal conditions, Sir2 levels were high enough that Sir2/3/4 and RENT complexes were not in conflict over Sir2. In contrast, *sir2-G436D* reduced Sir2-G436D levels such that it could not simultaneously meet the requirements of both the Sir2/3/4 and RENT complexes.

Though heterochromatic silencing is often framed as an epigenetic mechanism, our data suggested that genetically heritable differences in rDNA copy number is an additional mechanism that can lead to variable yet heritable expression states of heterochromatin. The genetic heritability of different rDNA copy numbers is broadly conserved (Lyckegaard and Clark 1989; Zhang *et al*. 1990; Gibbons *et al*. 2015), and it is interesting to speculate how cells either utilize or mitigate the effects of this variation. In yeast, different rDNA copy numbers are linked to differences in gene silencing, the monitoring of replication initiation, and replicative lifespan (Kaeberlein *et al*. 1999; Michel *et al*. 2005; Ganley *et al*. 2009). Whether these differences provide adaptive benefits or simply reflect the competition of different cellular processes over limiting factors, such as Sir2, will certainly be a motivating question for future studies.

### The existence of an intermediate silenced state

Single-cell analysis is useful to study heritable expression states in a cell population; this concept has been illustrated by multiple studies that uncovered and characterized the epigenetic states seen in *sir1*Δ (Pillus and Rine 1989; Xu *et al*. 2006). One important aspect of silencing in *sir1*Δ is that silenced cells are silenced to the same degree as *SIR+* cells, and expressed cells are expressed to the same degree as *sir2*Δ cells (Figure 2). In contrast, *sir2-G436D* exhibited a mix of silenced cells and cells that exhibited intermediate expression, as measured by flow cytometry and microscopy. Remarkably, these intermediate states were heritable through multiple cell divisions.

Curiously, overexpression of *sir2-G436D* did not influence the frequency of different expression states seen in *sir2-G436D* by flow cytometry, and *fob1*Δ had relatively small effects on this frequency. This result contrasted with the ability of *sir2-G436D* overexpression to partially reduce, and of *fob1*Δ to strongly reduce, variegation of silencing at the colony level. Together, these results suggested that the majority of switching events at the single cell level were independent of changes in rDNA copy number and the associated colony-level variegation. In this model, a relatively high switching rate between silencing states of *sir2-G436D* manifested as uniform, intermediate fluorescence at the colony level in *fob1*Δ. Then, the added layer of rDNA copy number changes in *FOB1* altered heritability of these silencing states in a manner that was relatively small at the single-cell level, but readily observed as radial streaks at the macroscopic level of a colony. Therefore, the intermediate expression state observed in *sir2-G436D* was mostly independent of changes in rDNA copy number and may have derived from a unique behavior of the Sir2-G436D protein at silenced loci.

A recent study found that Sir-based silencing establishment at both *HML* and *HMR* occurs through an intermediate silenced state, rather than an abrupt switch from the fully expressed to fully silenced state (Goodnight and Rine 2020). Furthermore, this intermediate state could be generated and stably maintained when certain histone modifying enzymes were absent in G1-arrested cells. Ultimately, that study concluded that silencing establishment occurs through a shift in the landscape of histone modifications at *HML* and *HMR*, and that cells that do not fully experience this shift can maintain a partially silenced state. In this view, the intermediate silencing state observed in *sir2-G436D* may reflect a partial deficiency in its ability to deacetylate H4K16. It is interesting to note that deletion of *SAS2*, which is responsible for acetylation of H4K16, also exhibits intermediate silencing states at *HML* and *HMR* at the single-cell level (Xu *et al*. 2006). Notably, the intermediate silencing state in *sas2*Δ is not a bona fide epigenetic state, as it is present in all cells of that genotype. Taken together, these results strongly suggest that defects in different histone modifying enzymes can exhibit similar phenotypes of intermediate silencing. This trend points to the existence of silencing intermediates that can be uncovered by modulating a complex landscape of histone modifications. Additional studies on *sir2-G436D*, *sas2*Δ, and other mutants will clarify how histone modifying complexes can shift the strength of silencing and, in some cases, reveal heritable properties of heterochromatin.

## Acknowledgements

We thank the members of our lab, especially Marc Fouet, for extensive discussions throughout the course of this work. This work was funded by a SURF L&S Fellowship through UC Berkeley (to DF), by an National Science Foundation fellowship (DGE1752814 to DSS), and by grants from the National Institutes of Health (GM31105 and GM120374, to JR).

## Data Availability

Strains and plasmids are available upon request. Supplemental files available at FigShare.

---

**Figure S1:**
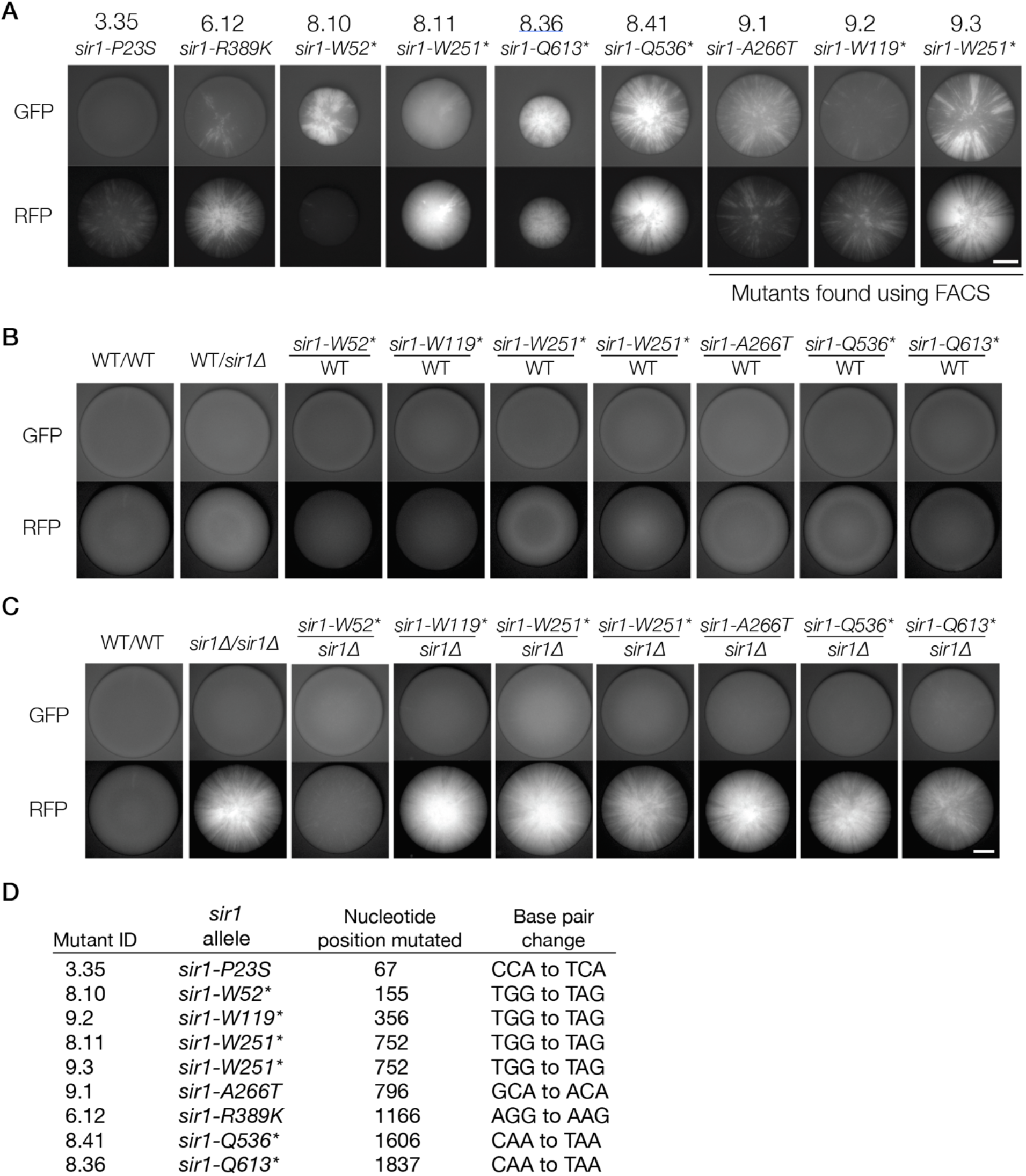
Additional FLAME phenotypes and characterization of the *sir1* mutants identified. **A.** Colony phenotypes of the mutants identified (JRY11896, JRY11897, JRY11901, JRY11902, JRY11904, JRY11905, JRY11918-11920). Colonies were labeled with both the initial mutant ID and its associated *sir1* allele; an asterisk indicates a premature stop codon. Each mutant found during FACs sorting was from an independent mutagenized culture. **B.** Colony images of the dominance test diploid strains (*MATa* FLAME strain crossed with a *MATα* mutant FLAME strain) (JRY11908, JRY11909, JRY11914-11917, JRY11922-11924, JRY11955), in both the GFP and RFP channel, plated on YPD. Some variable autofluorescence was seen in the RFP channel at the colony level, however variable autofluorescence in the RFP channel was also seen in colonies without an endogenous source of RFP (data not shown). **C.** Colony images of the *sir1*Δ complementation test diploid strains (*MATa sir1*Δ FLAME strain crossed with a *MATα* mutant FLAME strain) (JRY11946-11957), in both the GFP and RFP channel, plated on YPD. Mutant *sir1-W52\** displayed a much weaker phenotype than other colonies, yet some small sectors are seen in the RFP channel. **D.** Table indicating the initial mutant ID, the associated *sir1* allele, the location of the nucleotide mutation, and the resulting base pair change. Mutants 8.11 and 9.3 were isolated from independent rounds of mutagenesis and had identical *sir1* alleles, *sir-W251*\*. Scale bars, 2 mm.

**Figure S2:**
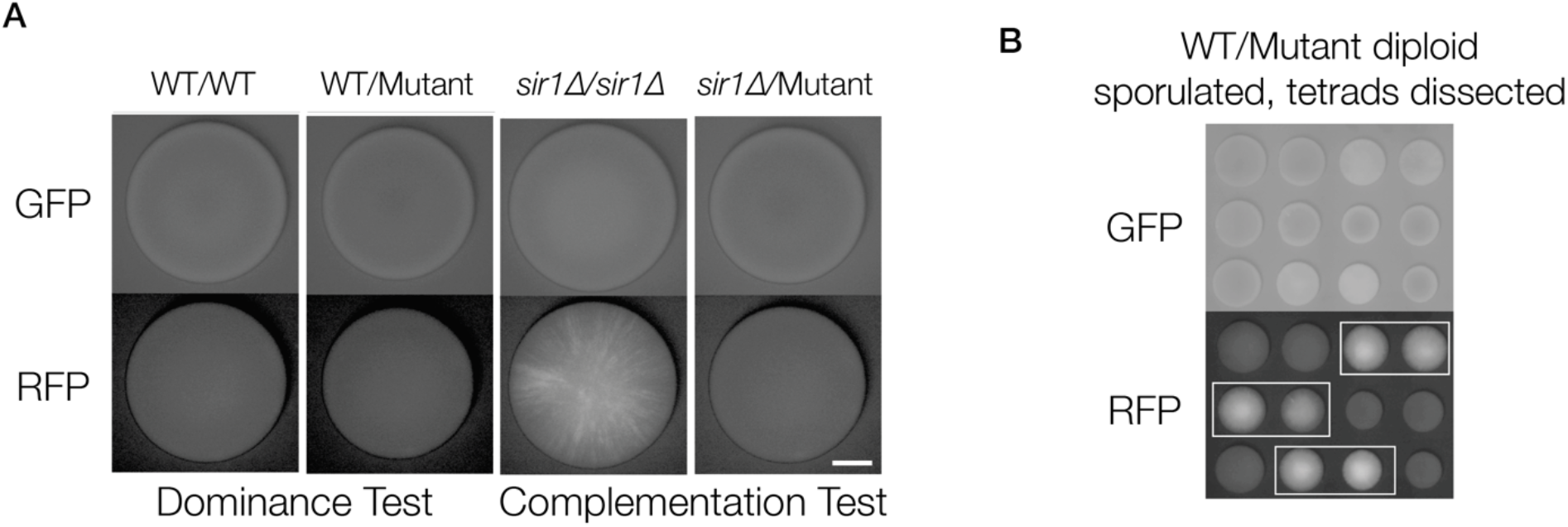
Genetic analysis of the mutant isolated from the second mutagenesis screen. A. Representative colony images of the diploid strains (JRY11957, JRY12476, JRY11952, JRY12477), grown on CSM. These strains were generated by mating the mutant isolated from the second mutagenesis screen (JRY12466) to *SIR+* and *sir1*Δ strains (JRY12863 and JRY12864, respectively). Scale bar, 2 mm. B. The wild-type/mutant diploid (JRY12476) was sporulated and tetrads were dissected on YPD.

**Figure S3:**
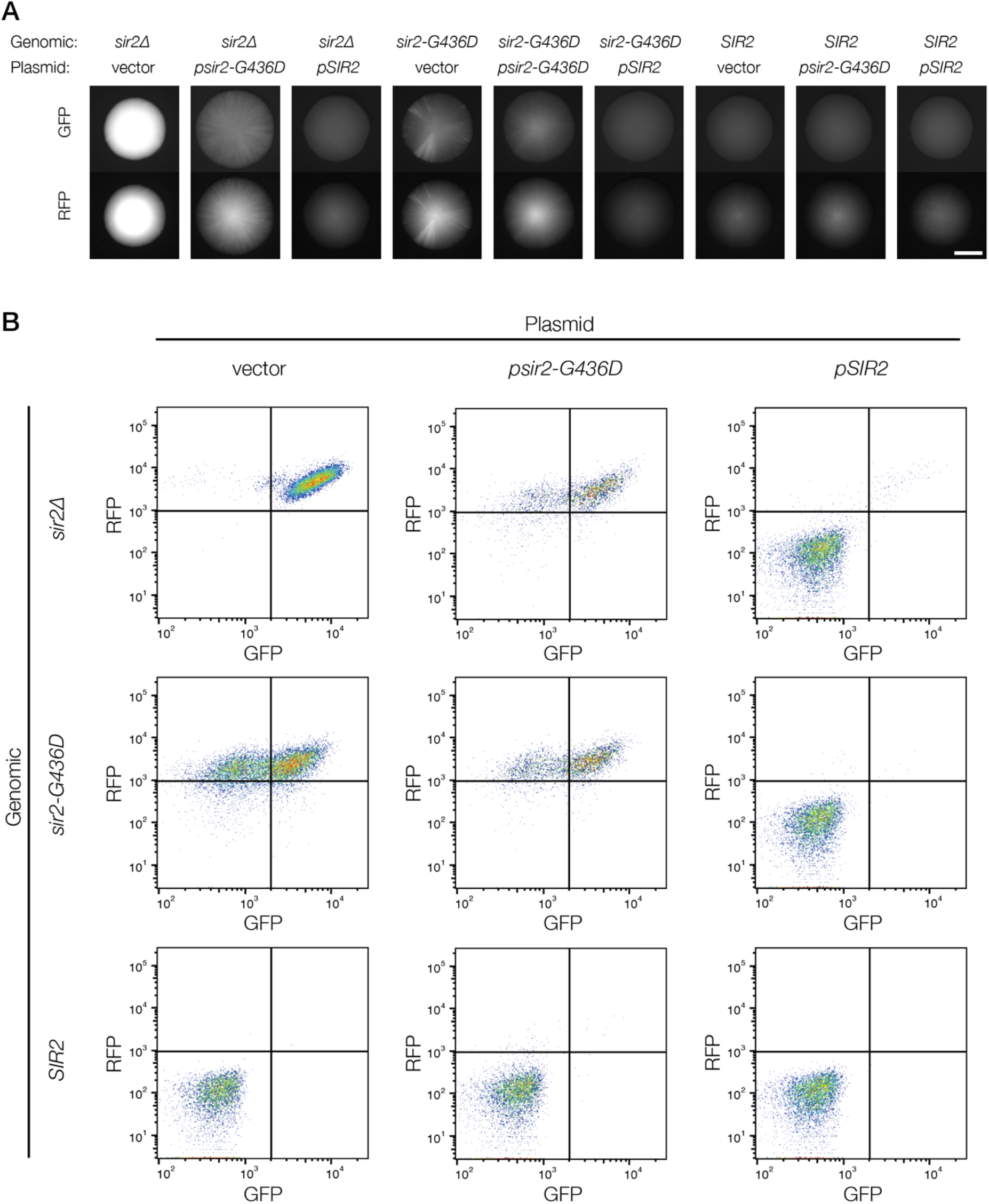
Effects of *SIR2* and *sir2-G436D* overexpression. A. Representative colony images of *sir2*Δ (JRY12259), *sir2-G436D* (JRY12564), and *SIR2* (JRY12860) strains with a 2 micron vector (pRS426), a 2 micron vector containing *sir2-G436D* (*psir2-G436D*) (pJR3525), or a 2 micron vector containing *SIR2* (*pSIR2*) (pJR3524). Colonies were grown on CSM −Ura to select for plasmids. Scale bar, 3 mm. B. Flow cytometry profiles of strains shown in (A). Independent cultures (n = 3 per genotype) were grown at log phase for 24 hours in CSM −Ura liquid media, fixed, and analyzed. Representative flow cytometry profiles for each strain are shown. Quadrants were established by using the fluorescence profiles of *SIR2* and *sir2*Δ cells.

**Figure S4:**
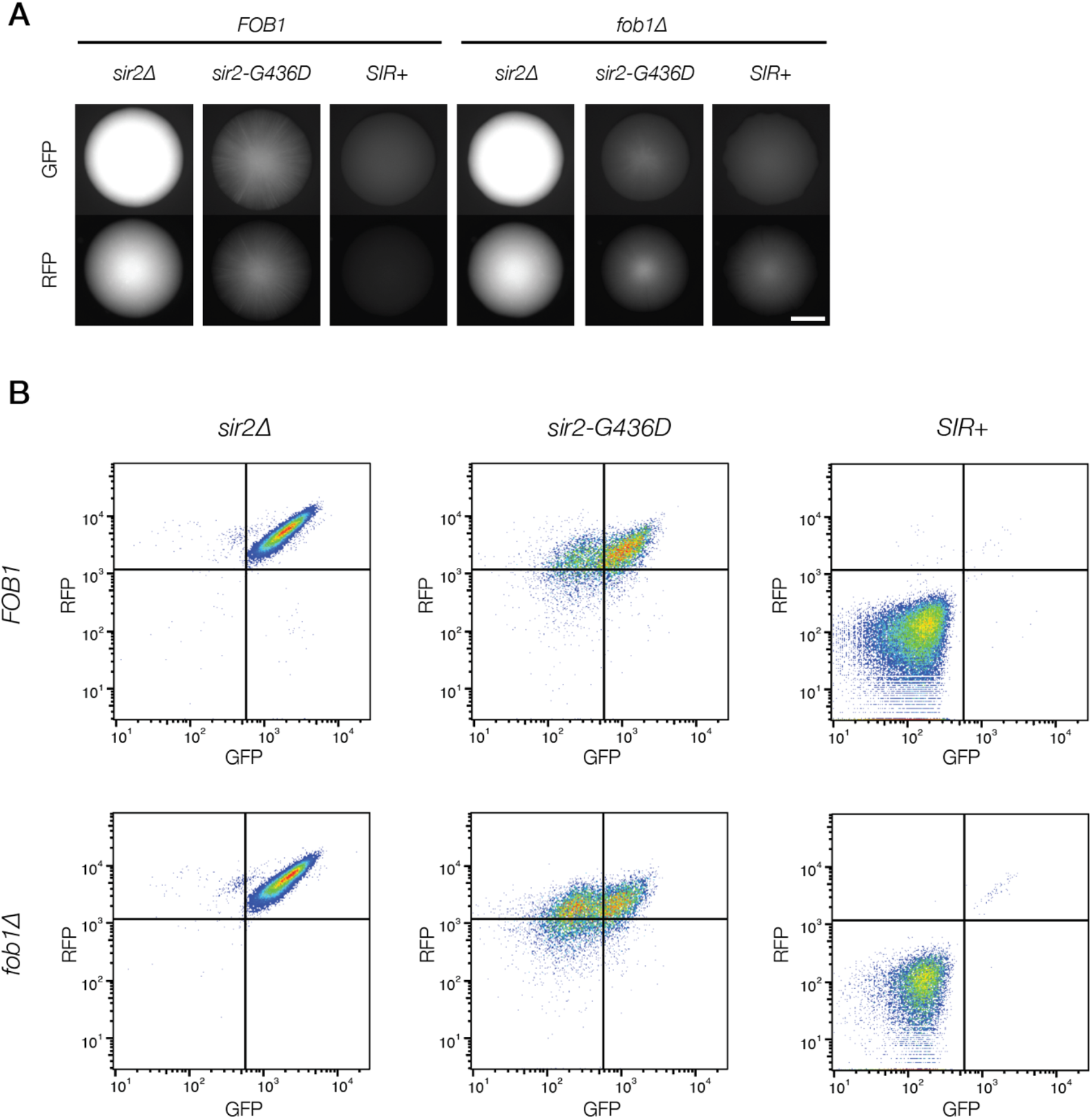
Effects of rDNA recombination on *sir2-G436D*. A. Representative colony images of *sir2Δ, sir2-G436D*, and *SIR2* with or without *FOB1* (JRY12259, JRY12564, JRY12860, JRY12900-12902), grown on CSM. Scale bar, 3 mm. B. Flow cytometry profiles of strains shown in (A). Independent cultures (n = 3 per genotype) were grown at log phase in CSM liquid media for 24 hours, fixed, and analyzed. Representative flow cytometry profiles are shown for each strain. Quadrants were established by using the fluorescence profiles of *SIR2* and *sir2*Δ cells.

**Supplemental Movie 1:** *sir1*Δ cells exhibited heritable epigenetic states at *HMR*. Cells were grown on CSM solid medium and monitored by fluorescence microscopy (JRY12861). Images represent overlays between bright-field and GFP channels. Scale bar, 10μm.

**Supplemental Movie 2:** *sir2-G436D* cells exhibited heritable epigenetic states at *HMR*. Cells were grown on CSM solid medium and monitored by fluorescence microscopy (JRY12564). Images represent overlays between bright-field and GFP channels. Scale bar, 10μm.

**Supplemental Movie 3:** *sir2-G436D, fob1*Δ cells exhibited heritable epigenetic states at *HMR*. Cells were grown on CSM solid medium and monitored by fluorescence microscopy (JRY12901). Images represent overlays between bright-field and GFP channels. Scale bar, 10μm.

